# White matter connectivity in uncinate fasciculus accounts for visual attention span in developmental dyslexia

**DOI:** 10.1101/2022.07.09.499403

**Authors:** Jingjing Zhao, Zujun Song, Yueye Zhao, Michel Thiebaut de Schotten, Irene Altarelli, Franck Ramus

## Abstract

The present study aimed to investigate the role of connectivity disruptions in two fiber pathways, the uncinate fasciculus (UF) and the frontal aslant tract (FAT), in developmental dyslexia and determine the relationship between the connectivity of these pathways and behavioral performance in children with dyslexia. A total of 26 French children with dyslexia and 31 age-matched control children were included. Spherical deconvolution tractography was used to reconstruct the two fiber pathways. Hindrance-modulated oriented anisotropy (HMOA) was used to measure the connectivity of each fiber pathway in both hemispheres. The boys with dyslexia showed reduced HMOA in the UF compared to control boys, but this difference was not observed in girls. Furthermore, HMOA of the UF correlated with individual differences in the visual attention span in participants with dyslexia. All significant results found in HMOA of the UF were verified in fractional anisotropy (FA) of the UF using standard diffusion imaging model. This study suggests a differential sex effect on the connectivity disruption in the UF in developmental dyslexia. It also indicates that the UF may play an essential role in the visual attention span deficit in developmental dyslexia.

**Significance Statement:** This study presents the first account of connectivity disruption in the uncinate fasciculus in developmental dyslexia. In particular, this connectivity disruption only appears in boys with dyslexia but not in girls with dyslexia. We also show that the connectivity of the uncinate fasciculus accounts for individual differences in the visual attention span in children with dyslexia, expanding the current understanding of the function of the uncinate fasciculus.

## Introduction

Developmental dyslexia (DD) is defined as a developmental learning disorder with impairment in reading, which is not due to a disorder of intellectual development, sensory impairment (vision or hearing), neurological or motor disorder, lack of availability of education, lack of proficiency in the language of academic instruction, or psychosocial adversity (World Health Organization, 2018). The prevalence of DD ranges from 1.3 to 17.5 % (Di Folco et al., 2020; Shaywitz and Shaywitz, 2005).

DD has been widely accepted as a neurodevelopmental learning disorder that manifests as a neural disconnection syndrome (Boets et al., 2013; Horwitz et al., 1998; Paulesu et al., 1996; Pugh et al., 2001; Zhao et al., 2016; Lou et al., 2019; Liu et al., 2021, 2022). Studies using diffusion tensor imaging (DTI) techniques have identified DD-related anomalies in white matter pathways such as the arcuate fasciculus (AF), superior longitudinal fasciculus (SLF), inferior frontal-occipital fasciculus (IFOF), and inferior longitudinal fasciculus (ILF) (Carter et al., 2009; Keller & Just, 2009; Odegard et al., 2009; Richards et al., 2008; Rimrodt et al., 2010; Vandermosten et al., 2012; Yeatman et al., 2012; Zhao et al., 2016; Su et al., 2018). These white matter pathways mainly correspond to two reading networks: the dorsal network (AF and SLF) and the ventral network (IFOF and ILF) (Hickok & Poeppel, 2007; Vandermosten et al., 2012; Frühholz et al., 2015; Oliver et al., 2017; Rollans and Cummine, 2018; Su et al., 2018; Yablonski et al., 2019; Bhattacharjee et al., 2020). However, prior studies never examined two white matter pathways that are potentially important for reading and DD: the uncinate fasciculus (UF) and frontal aslant tract (FAT). Like the AF and the IFOF, these two white matter pathways connect with the inferior frontal lobe, including Broca’s area.

The uncinate fasciculus (UF) is one of the white matter structures associated with the ventral reading network (Schlaggar and McCandliss, 2007), which connects the anterior region of the temporal lobe to the frontal lobe (Catani et al., 2002). The UF has been found to play an essential role in language, cognitive, and social abilities (Papagno et al., 2011; Weis et al., 2018). Prior studies have shown that the function of the UF is related to semantic processing, such as word understanding and selecting the appropriate semantic representation from a set of activation representations in semantic memory (Harvey et al., 2013; Di Tella et al., 2020). It has been demonstrated that the connectivity of the left UF was highly correlated with the performance of semantic processing scores in primary progressive aphasia (PPA) (Catani et al., 2013; Harvey et al., 2013). In addition, a disruption in the UF has also been commonly found to be associated with anomalies of social behaviors in neuropsychiatric disorders, including autism spectrum disorder (ASD) (Kumar et al.,2010; Pugliese et al., 2009), psychopathy (Craig et al., 2009; Sundram et al., 2012), and social anxiety disorders (Baur et al., 2013; Phan et al., 2009).

The frontal aslant tract (FAT) was discovered by Catani and colleagues, who defined it as a fiber pathway connecting the middle frontal lobe (SMA and pre-SMA) and Broca’s region (Lawes et al., 2008; Oishi et al., 2008; Catani et al., 2012; Thiebaut de Schotten et al., 2012). Recently, the FAT was found to play an essential role in the production of language (Catani et al., 2012; Catani et al., 2013; Dick et al. 2014; Vassal et al., 2014). For example, it has been demonstrated that connectivity anomalies of the left FAT were associated with verbal fluency defects in PPA (Catani et al., 2013).

In sum, although some studies have demonstrated a link between disruptions of the UF and FAT and impairment in neurological disorders, no study to date has examined whether these two fiber pathways show disruptions in DD, nor have they explored the function of these two fiber pathways in DD. Therefore, the aim of this study was twofold. First, we aimed to determine whether there were connectivity disruptions in the UF and FAT in DD. Second, if any disruption was observed, we aimed to reveal further whether the connectivity of the fiber pathways could account for individual differences in cognitive and reading-related skills in DD. Finally, the uneven sex ratio in dyslexia in favor of males has been known for a long time (Galaburda et al., 1985 a b c; Rutter et al., 2004), and it has been suggested that the neural basis of dyslexia might partly differ between males and females (Ramus et al., 2018). Therefore, we will also investigate sex as a factor and a potential moderator of group differences.

## Method

### Research transparency and Data availability

The experimental procedures and analyses of the uncinate fasciculus were preregistered on the Open Science Framework (OSF, http://osf.io/uxk8a). Further details can also be found in our previously published paper which used the same dataset (Zhao et al., 2016).

The conditions of our ethics approval do not permit public archiving of anonymized study data. Readers seeking access to the data should contact Franck Ramus and Jingjing Zhao.

### Participants

A total of 26 children with dyslexia and 31 control children (aged 9-14 years) matched in sex, age, handedness, and nonverbal IQ were included in this study (see participant characteristics in Table 1). All children were native French speakers with normal vision and hearing. No child had a history of neurological or psychiatric disorders. Based on the Alouette test, which assesses reading accuracy and speed (Lefavrais, 1967), children with dyslexia were included based on a delay larger than 18 months on text reading age, while children were assigned to the control group if they were no more than 12 months behind. The study was approved by the ethics committee of Bicêtre Hospital, and informed consent was obtained from all children and their parents.

**Table 1.**
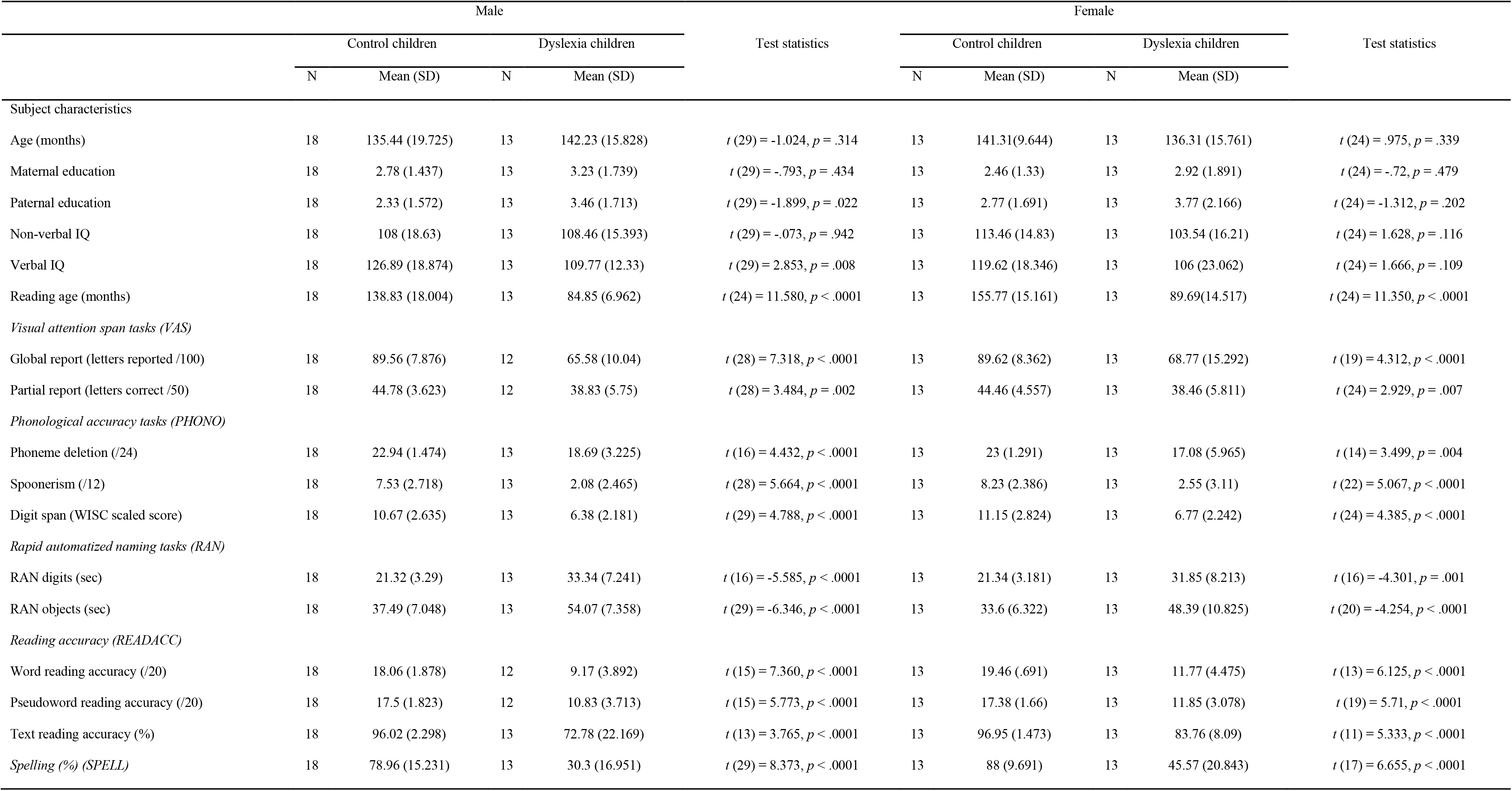
Demographical data and behavioral results of reading-related abilities.

### Behavioral measures

A series of behavioral tests were used to assess the cognitive and literacy skills of each child. The intellectual abilities of the children were assessed by WISC blocks, matrices, similarities, and comprehension subtests (Wechsler, 2005). Their visual attention span was measured by both the global and the partial letter report tasks (Bosse et al., 2007; Saksida et al., 2016). Phonological awareness was tested by a phoneme deletion task (Sprenger-Charolles et al., 2005) and a spoonerism test (Bosse and Valdois, 2009). Verbal working memory was tested with the WISC digit span subtest (Wechsler, 2005). Phonological processing speed was estimated by rapid automatized naming (RAN) tasks for digits and objects (Plaza and Robert-Jahier, 2006). Text reading fluency was assessed by the Alouette test (Lefavrais, 1967). The Odedys test was used to estimate word and nonword reading ability (Jacquier-Roux et al., 2005). Orthographic skill was assessed by a word spelling-to-dictation test (Martinet and Valdois, 1999).

For further correlation analysis with brain measures, we calculated average z-scores to define five composite measures as follows: visual attention span (VAS) from global and partial letter report; phonological processing abilities (PHONO) from phoneme deletion, spoonerisms, and digit span; RAN from digit and object RAN; reading accuracy (READACC) from words, pseudowords, and text reading accuracy; spelling (SPELL) was simply the z-score of the word spelling-to-dictation test. Signs were adjusted such that positive z-scores represented above-average performance.

### Image Acquisition and Analysis

A 3T MRI scanner (Tim Trio, Siemens Medical Systems, Erlangen, Germany) was equipped with a whole-body gradient (40 m T/m, 200 T/m/sec) and a 32-channel head coil to scan all the children. A MPRAGE sequence with the parameters TR=2,300 ms; TE=3.05 ms; flip angle = 9^°^; field of view =230 ms; voxel size = 0.9 × 0.9 × 0.9 mm^3^; acquisition matrix= 230 × 230 × 224 was used to acquire T1-weighted structural MRI scans. A spin-echo single-shot EPI sequence was used for diffusion MRI scans, with parallel imaging (GRAPPA reduction factor 2), partial Fourier sampling (factor 6/8), and bipolar diffusion gradients to reduce geometric distortions. The whole brain was imaged with an isotropic spatial resolution of 1.7 mm^3^ (matrix size = 128 × 128, a field of view = 218 mm) and 70 interleaved axial slices. Diffusion gradients were applied along 60 orientations, uniformly distributed, with a diffusion weighting of b = 1400 sec/mm^2^ (repetition time = 14,000 msec, echo time = 91 msec). Additionally, three images were acquired with no diffusion gradient applied (b = 0). Each sequence took about 6 min, resulting in a total acquisition time of 18 min.

We first integrated the three sequences of raw data into a single data file. Then, we used ExploreDTI (Leemans and Jones, 2009; http://www.exploredti.com) to simultaneously register and correct motion and geometrical distortions in images of subjects. We adopted a damped Richardson-Lucy algorithm for spherical deconvolution (SD) (Dell’Acqua et al., 2010) to estimate multiple orientations in voxels containing different populations of crossing fibers for our high angular resolution diffusion-weight imaging (HARDI) data. Algorithm parameters were chosen following Zhao et al. (2016) and Thiebaut de Schotten et al. (2011).

At least one fiber orientation was selected as a seed in every brain voxel to establish whole-brain tractography. For each fiber orientation in these voxels, Euler integration was used, with a step size of 1 mm, to propagate streamlines (Dell’Acqua, Simmons, Williams, & Catani, 2013). The algorithm followed the orientation vector of least curvature (Schmahmann et al., 2007) when entering a region that crossed white matter fibers. Streamlines ended when a voxel without fiber orientation was reached, or the curvature between two steps exceeded a threshold of 60°. Spherical deconvolution, fiber orientation vector estimations, and tractography were performed using in-house software developed with MATLAB v7.8 (The MathWorks, Natick, MA).

Tract dissections were performed in the native space using TrackVis for each child (http://trackvis.org/, see Wedeen et al., 2008), which allows for the identification of the tracts, visualization in 3 dimensions, and quantitative analyses on each fiber. The region-of-interest (ROI) approach was adopted to extract the tracts of interest, and the principle for defining the ROIs for each fiber tract was based on previous tractography studies on the uncinate fasciculus (Catani et al., 2002; Catani et al., 2013; Harvey et al., 2013; Di Tella et al., 2020) and the frontal aslant tract (Catani et al., 2013; Thiebaut de Schotten et al., 2012).

To limit inter-subject variability related to the operator expertise and automate some tract dissection steps, the FMRIB Software Library package (https://fsl.fmrib.ox.ac.uk/fsl/fslwiki/) was used to define ROIs on the MNI152 template. A contrast map for white matter named a convergence map (CS maps; Dell’Acqua et al., 2006) was calculated using the Richardson-Lucy Spherical Deconvolution Algorithm for each subject. Then, the convergence map was registered to the MNI152 template by Advanced Normalization Tools (http://picsl.upenn.edu/software/ants/), combining affine with diffeomorphic deformations (Avants et al., 2008; Klein et al., 2009). The inverse deformation was then applied to the ROIs defined on the MNI152 template to bring them into the native space of every participant.

The individual dissections of tracts in the native brain space of each participant were then visually inspected and corrected by four anatomists (JZ, MTS, ZS, and YZ). UF and FAT of all participants were successfully reconstructed in the HARDI model using spherical deconvolution tractography. Hindrance-modulated oriented anisotropy (i.e., HMOA; Dell’Acqua et al., 2013) was extracted on every dissected pathway, used as a compact measure of fiber density and connectivity characterizing the diffusion properties along each track orientation. The average HMOA across each entire tract was regarded as the main variable of interest. The validation analysis of spherical deconvolution tractography in dyslexia research has been reported in a previous study (Zhao et al., 2016).

Because most prior studies on the white matter connectivity of the uncinate fasciculus and the frontal aslant tract adopted the standard diffusion tensor imaging (DTI) model (Catani et al., 2013; Harvey et al., 2013), we also used the standard DTI model to verify the analyses based on HARDI model, to test the robustness of our results. The whole-brain tractography was imported to Trackvis, and the definition of ROIs in the HMOA analysis was used for the UF and FAT in Trackvis. The mean fractional anisotropy (FA) value was computed for both hemispheres. The UF of 7 participants could not be reconstructed in the standard DTI model: one bilaterally and six in the left hemisphere. The FAT could not be reliably reconstructed using the standard DTI model in the left and right hemispheres in about half of all participants. Because the sample size of the FAT that was successfully reconstructed using the standard DTI model was too small, we only performed FA analysis for the UF.

### Statistical analysis

Statistical analysis was performed using SPSS software (SPSS26, Chicago, IL). An independent t-test or Chi-squared test was adopted to measure behavioral and demographic differences between control subjects and participants with dyslexia. With regard to the microstructure of the white matter pathways, general linear models with repeated measures were run separately for the UF and FAT, with the mean HMOA measure of each tract as a dependent variable, group (control vs. dyslexia), and sex (male vs. female) as between-subject variables, and hemisphere as a within-subject variable. Age and parental education were used as covariates in the model. Post-hoc comparisons further investigated significant interactions between groups and other factors.

Pearson partial correlation analysis was used to examine the correlation between the HMOA of the white matter pathways that showed significant group differences and cognitive and literacy skills, controlling for the effects of age, sex, and parental education level. The correlation results were Bonferroni corrected for two groups and five behavioral tests (0.05/10=0.005). Significant effects were further confirmed by analyzing FA using the standard DTI model.

Finally, since significant correlations were observed between visual attention span and HMOA and FA of the UF in dyslexic individuals, and visual attention span and phonological deficits usually show comorbidity in dyslexia (Liu et al., 2022; Peyrin et al., 2012), hierarchical liner regression models were further conducted. The regression models used HMOA and FA of UF as dependent variables, and visual attention span as independent variables, controlling for age, sex, parental education in the first step, and phonological processing abilities (PHONO: phoneme deletion, spoonerisms, and digit span) in the second step.

## Results

### Demographics and behavioral results

Demographic and behavioral measures for the control and dyslexia groups are shown in Table 1. There were no group differences for age, sex, handedness, or non-verbal IQ. The two groups were also matched in maternal educational level. Individuals with dyslexia performed worse than control children for all phonological and literacy skills.

### White matter connectivity analysis HMOA analysis

The three-way ANOVA revealed no main effects of group (F _(1,51)_ =0.591, *p* = 0.446, *η_p_^2^*= 0.011) or hemisphere (F _(1,51)_ =2.585, *p* = 0.114, *η_p_^2^* = 0.048) in HMOA of the UF. A trend of main effect of sex (F _(1,51)_ =3.729, *p* = 0.059, *η_p_^2^*= 0.068) was observed: HOMA of the UF in boys was lower than that in girls. However, the interaction between group and sex in HMOA of the UF was significant (F _(1,51)_ =7.036, *p* = 0.011, *η_p_^2^*= 0.121, see Figure 1, Table 2). Post-hoc analysis showed the boys with dyslexia had a reduction in HMOA in the UF compared with control boys (Control: *Mean ± SD* =0.098 ± 0.0066, Dyslexia: *Mean ± SD* =0.091 ± 0.0096, *p*=0.016, Cohen’s d= 0.819), but not in girls (Control: *Mean ± SD* =0.097 ± 0.0066, Dyslexia: *Mean ± SD*=0.099 ± 0.007, *p*=0.208, Cohen’s d=0.368). No other significant results for interaction effects between group and other variables in UF were found. No significant effects in the HMOA of the FAT were observed.

**Figure 1.**
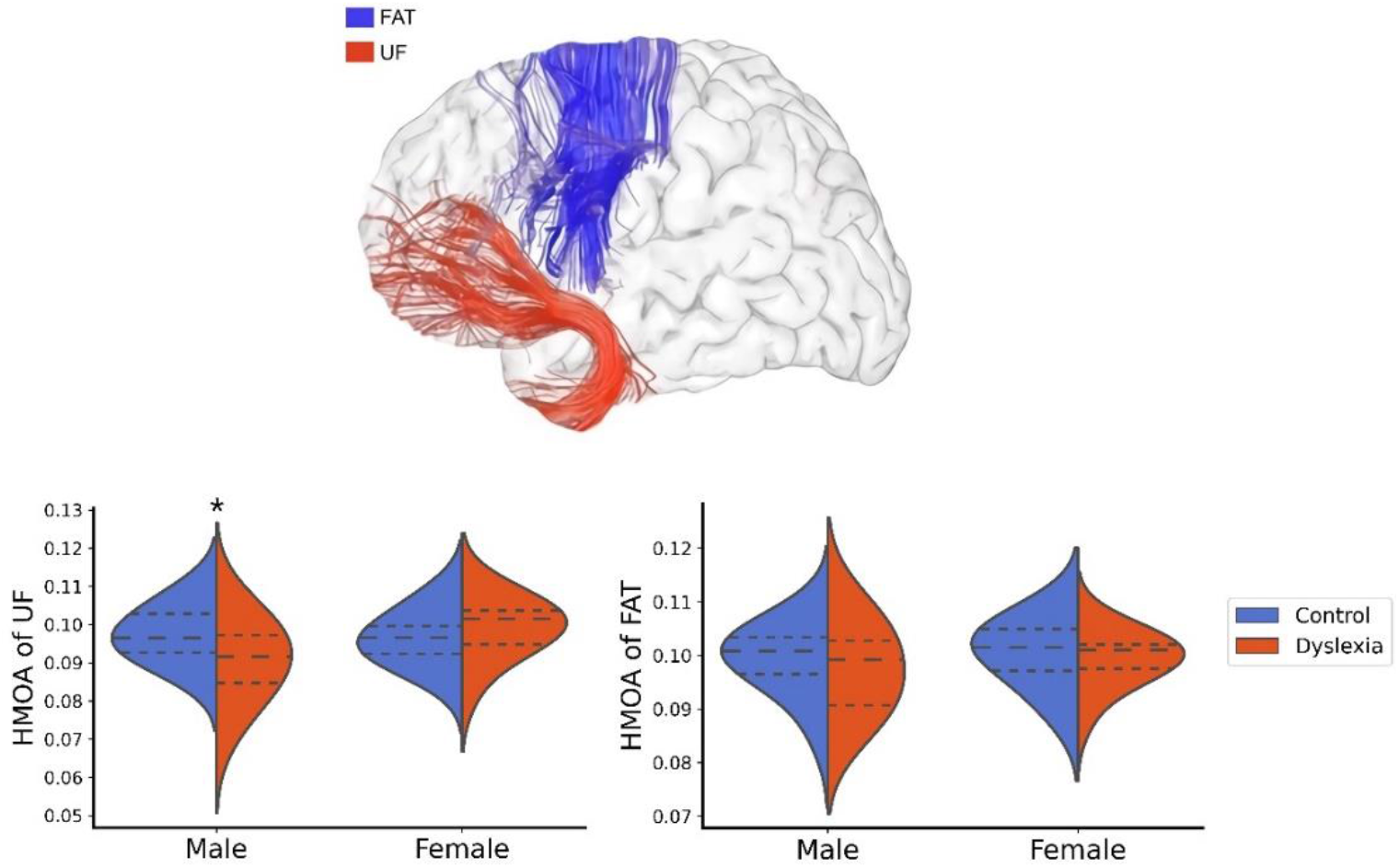
Differences in hindrance-modulated oriented anisotropy (HMOA) in uncinate fasciculus (UF) and frontal aslant tract (FAT) between dyslexia and control groups in males and females respectively. **\****p* < 0.05

**Table 2.** Mean hindrance-modulated oriented anisotropy (HMOA) of uncinate fasciculus (UF) and frontal aslant tract (FAT) in control and children with dyslexia.

|  |  | Male |  |  |  | Female |  |  |  |
| --- | --- | --- | --- | --- | --- | --- | --- | --- | --- |
|  |  | Control children |  | Dyslexia children |  | Control children |  | Dyslexia children |  |
|  |  | N | Mean (SD) | N | Mean (SD) | N | Mean (SD) | N | Mean (SD) |
| UF_HMOA | left | 18 | 0.0972 (0.0090) | 13 | 0.0889 (0.0101) | 13 | 0.0965 (0.0080) | 13 | 0.0987 (0.0091) |
|  | right | 18 | 0.0979 (0.0077) | 13 | 0.0927 (0.0109) | 13 | 0.0968 (0.0070) | 13 | 0.0995 (0.0063) |
| FAT_HMOA | left | 18 | 0.0996 (0.0055) | 13 | 0.0992 (0.0075) | 13 | 0.1024 (0.0060) | 13 | 0.1006 (0.0058) |
|  | right | 18 | 0.0988 (0.0077) | 13 | 0.0960 (0.0083) | 13 | 0.0984 (0.0072) | 13 | 0.0980 (0.0065) |

The HMOA of the UF showed a positive correlation with VAS in individuals with dyslexia (*r* = 0.584, *p* = 0.004, surviving Bonferroni correction, *p* < 0.05/10=0.005 R*^2^*= 0.341, see Table 3 and Figure 2). No significant correlations were observed between the HMOA of the UF and other behavioral measures in the children with dyslexia and the control group. Hierarchical liner regression analysis (Table 4) of the HMOA of UF showed that VAS remained a significant predictor of the HMOA of UF after controlling age, sex, parental education, and PHONO (*β* = 0.559, *p* = 0.004).

**Figure 2.**
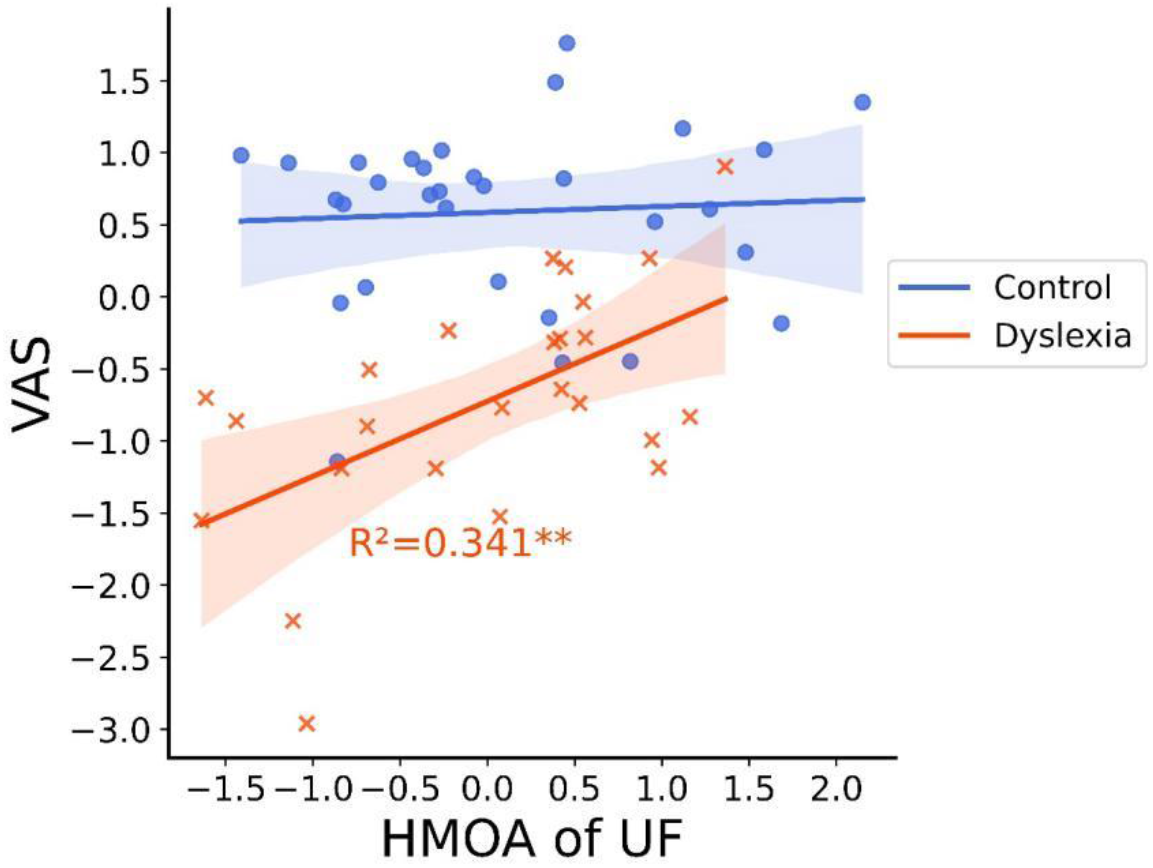
Partial correlation (covariate: sex, age, parental education) between microstructure of the uncinate fasciculus (UF) and visual attention span (VAS) in control and children with dyslexia respectively. \*\**p* < 0.01

**Table 3.** Partial correlation coefficients (controlled for age, sex, and parental education) between hindrance-modulated oriented anisotropy (HMOA) of uncinate fasciculus (UF) and behavioral measures of visual attention span (VAS), phonological processing abilities (PHONO), rapid automatized naming (RAN), reading accuracy (READACC), and spelling (SPELL). \**p* < 0.05, \*\**p* < 0.01, \*\*\**p* < 0.005 (surviving Bonferroni correction).

|  |  |  | VAS | PHONO | RAN | READACC | SPELL |
| --- | --- | --- | --- | --- | --- | --- | --- |
| UF_HMOA | Control | r | 0.048 | 0.169 | 0.043 | -0.222 | -0.04 |
|  |  | p | 0.807 | 0.39 | 0.827 | 0.256 | 0.841 |
|  | Dyslexia | r | 0.584*** | 0.141 | 0.009 | 0.009 | -0.039 |
|  |  | p | 0.004 | 0.522 | 0.966 | 0.969 | 0.859 |

**Table 4.**
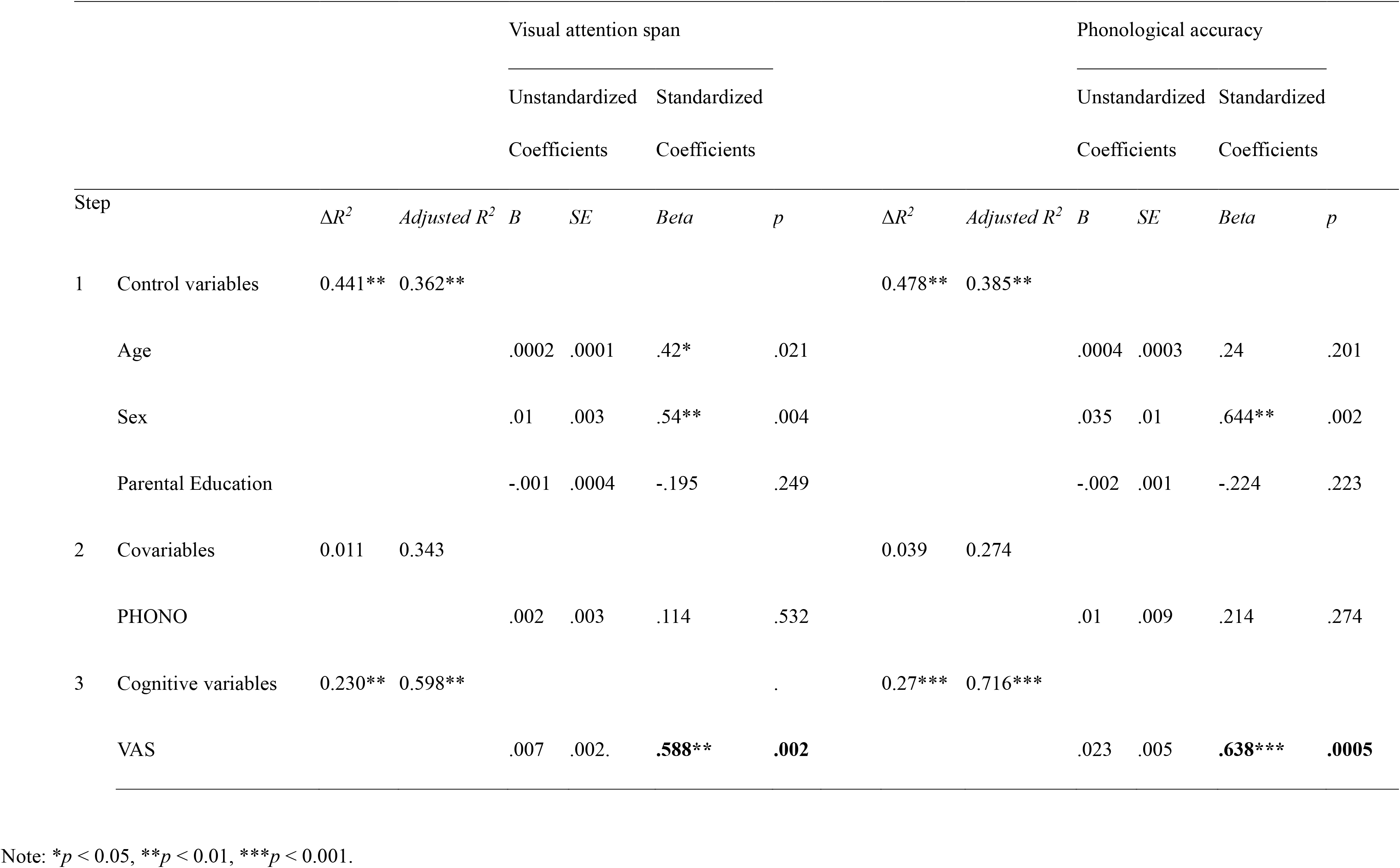
Hierarchical regression models using control variables and cognitive skills to predict the hindrance-modulated oriented anisotropy (HMOA) of uncinate fasciculus (UF).

### Verification FA analysis of UF

Correlations between FA and HMOA of UF were higher than 0.5 in both left hemisphere (*r* = 0.796, *p* < 0.001) and right hemisphere (*r* = 0.776, *p* < 0.001). Statistical analyses revealed a significant interaction between group and sex in the FA of UF (F _(1,44)_ =6.080, *p* = 0.018< 0.05, *η_p_^2^*= 0.121, Figure 3A). Specifically, the children with dyslexia showed a reduction in FA in the UF compared with controls; this reduction appeared only in boys (Control: *Mean ± SD* =0.42 ± 0.014, Dyslexia: *Mean ± SD* =0.403 ± 0.021, *p*=0.026, Cohen’s d=0.966), and not in girls (Control: *Mean ± SD* =0.429 ± 0.013, Dyslexia: *Mean ± SD* =0.435 ± 0.024, *p*=0.246, Cohen’s d=0.354). In addition, the FA of the UF showed a positive correlation with VAS in participants with dyslexia (*r* = 0.590, *p* = 0.01, R*^2^*= 0.348, see Figure 3B). Hierarchical liner regression analysis (Table 4) of the FA of UF also showed VAS remained a significant predictor of the FA of UF when control variables (age, sex, and parental education) and PHONO were statistically controlled (*β* = 0.638, *p* < 0.001). In short, the FA results based on the standard DTI model are consistent with the HMOA results using the HARDI model.

**Figure 3.**
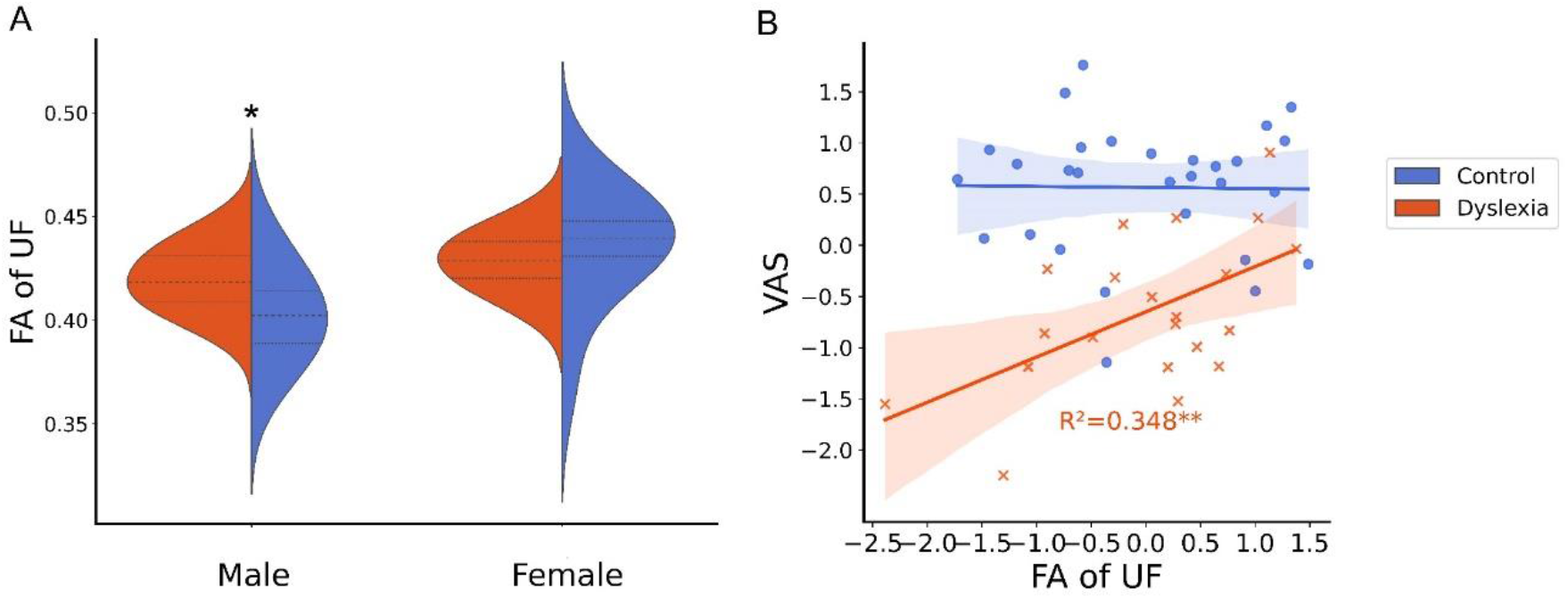
A: Differences in fraction anisotropy (FA) in uncinate fasciculus (UF) between dyslexia and control groups in males and females respectively; B: Partial correlation (covariates: sex, age, parental education) between microstructure of the uncinate fasciculus (UF) and visual attention span (VAS) in control and children with dyslexia respectively. *Note:* **\****p* < 0.05, \*\**p* < 0.01

## Discussion

The current study examined the connectivity disruption in the UF in children with dyslexia and explored the correlation between the microstructure of the UF and cognitive and reading abilities. It revealed a difference in the HMOA of the UF between participants with dyslexia and control children, but the difference was only found in boys and not in girls. The uncinate fasciculus also showed individual differences correlating with visual attention span in children with dyslexia.

First, this study revealed that a connectivity disruption in the UF is linked to developmental dyslexia. Indeed, prior studies have reported a connectivity disruption in the uncinate fasciculus in a range of developmental neurological diseases (Craig et al., 2009; Kumar et al., 2010; Catani et al., 2013; Harvey et al., 2013; Di Tella et al., 2020), but none of them have shown disruption of the UF in participants with dyslexia. Here, we provide first-hand evidence of connectivity disruption in the UF in developmental dyslexia.

Second, the connectivity disruption in the UF only appeared in boys with dyslexia and not in girls with dyslexia. This was the first time a differential sex effect was observed in a white matter connectivity disruption in children with dyslexia. Although several previous studies have reported sex differences in dyslexia, they only found brain structure differences in grey matter (Altarelli et al., 2013; Evans et al., 2014; Galaburda et al., 1985). Our results were consistent with these prior grey matter findings and provided new evidence that males with dyslexia exhibit atypical development of brain structure in white matter connectivity. Our results have therefore extended previous studies and provided further support to Galaburda and Geschwind (1985a, b, c)’s testosterone hypothesis, which was proposed to explain the higher incidence of dyslexia in males than in females – males are twice as likely to have dyslexia as females (Rutter et al., 2004). Progesterone and estrogen are believed to have protective effects on female cognition and nerves (Brann et al., 2007; Dumitriu., 2010), which implies that girls with dyslexia are less likely than boys with dyslexia to show physical changes at the level of brain structure compared to typically developing individuals. Although the above theoretical assumptions have not been fully proven, our research has contributed new evidence to our understanding of sex differences in developmental dyslexia and highlighted that sex is a potential factor for the heterogeneity of imaging research results in dyslexia (Ramus et al., 2018). However, it should be acknowledged that this study only recruited a limited number of children with dyslexia (13 boys and 13 girls), which might undermine the reliability of the findings of sex differences. Future studies might be valuable to further validate the current sex differences by including adequate sampling of the female and male dyslexic population.

Last but most importantly, our study revealed that the uncinate fasciculus exhibits individual differences that correlate with the performance of visual attention span in children with dyslexia. Moreover, our study revealed that the association between the uncinate fasciculus and visual attention span is independent of phonological processing. This finding is consistent with previous studies that have also reported the dissociated neural pattern when performing phonological and visual attention task (Peyrin et al., 2012; Liu et al., 2022). Nevertheless, the function of the left UF has been found associated with semantic processing in aphasic and brain-damaged individuals (Catani et al., 2013; Harvey et al., 2013; and Han et al., 2013). Bosse and Valdois (2009) found a sustained influence of visual attention span on reading performance, but only for irregular words, which involve more semantic processing. Therefore, the function of the UF in our individuals with dyslexia’s sample may tend to support a role in semantic processing rather than phonological processing. Yet, one limitation of the current study was that we did not adopt a behavioral task to test the semantic processing ability in children with dyslexia directly. Hence, we could not provide more evidence that the function of the UF is more related to visual attention span or semantic processing; this suggests a direction for future research.

Alternatively, prior studies have also indicated that the uncinate fasciculus is associated with the retrieval of word form and word production (Nomura et al., 2013; Papagno et al., 2011), rapid visual learning, which might be associated with the acquisition of language and attention shifts (Thomas et al., 2015; Kristjánsson, 2006), and the accuracy of stimulus judgment when executing high attention resource tasks (Di Tella et al., 2020). Thus, we cannot rule out the possibility that the function of the UF in our sample of dyslexia may also be related to orthographic processing and visual-attentional processes. At any rate, the current results suggest that the UF plays an essential role in the visual attention span in developmental dyslexia, and it may account for individual differences in visual attention span deficit in developmental dyslexia. In this way, we provide a new perspective and increase the understanding of the cognitive function of the uncinate fasciculus in dyslexia.

Finally, it should be noted that prior studies found dysfunction in the bilateral superior parietal lobules in individuals with dyslexia when performing visual attention span tasks (Lobier et al., 2014; Peyrin et al., 2012; Valdois et al., 2019). Here, we showed for the first-time evidence that the uncinate fasciculus may also account for the visual attention span deficit in dyslexia. We speculate, therefore, that there might be two neural networks that are both associated with visual attention span in dyslexia: a dorsal network (bilateral superior parietal lobe) related more to visual processing and a ventral network (uncinate fasciculus) correlated more with semantic processing.

## Conclusion

Our study reports a connectivity disruption in the uncinate fasciculus in developmental dyslexia. The connectivity disruption in the uncinate fasciculus was mainly manifested in boys with dyslexia but not in girls with dyslexia. The function of the uncinate fasciculus might be associated with visual attention span in developmental dyslexia.

## Authorship contribution statement

**Jingjing Zhao:** Conceptualization, Formal analysis, Methodology, Writing – original draft, Writing – review & editing, Supervision, Project administration, Validation, Investigation, Funding acquisition, Project administration. **Zujun Song:** Formal analysis, Writing – original draft, Writing – review & editing, Validation, Visualization. **Yueye Zhao:** Formal analysis, Validation, Visualization. **Michel Thiebaut de Schotten:** Conceptualization, Methodology, Software, Supervision, Writing – review & editing, Formal analysis. **Irene Altarelli:** Investigation. **Franck Ramus:** Resources, Supervision, Investigation, Methodology, Project administration, Writing – review & editing, Funding acquisition.

## Acknowledgments

The study was funded by National Natural Science Foundation of China (61807023), Humanities and Social Science Fund of Ministry of Education of the People’s Republic of China (17XJC190010), Shaanxi Province Natural Science Foundation (2018JQ8015), and Fundamental Research Funds for the Central Universities (CN) (GK201702011) to JZ. The study was also funded by Agence Nationale de la Recherche (ANR-06-NEURO-019-01, ANR-11-BSV4-014-01, ANR-17-EURE-0017 and ANR-10-IDEX-0001-02), Commission of the European Communities (LSHM-CT-2005-018696), École des Neurosciences de Paris Île-de-France to FR. MTS received funding from the European Research Council (ERC) under the European Union’s Horizon 2020 research and innovation programme (grant agreement no. 818521). The authors would like to thank all children who participated in the study and their parents, for their time and cooperation. We acknowledge the collaboration of Catherine Billard, Stéphanie Iannuzzi, Nadège Villiermet, Ghislaine Dehaene-Lambertz and all the staff at Neurospin for recruitment and testing, and we thank Sylviane Valdois for providing the visual attention span tests.

